# GeoAB: Towards Realistic Antibody Design and Reliable Affinity Maturation

**DOI:** 10.1101/2024.05.15.594274

**Authors:** Haitao Lin, Lirong Wu, Yufei Huang, Yunfan Liu, Odin Zhang, Yuanqing Zhou, Rui Sun, Stan Z. Li

**Affiliations:** Zhejiang University; AI Lab, Research Center for Industries of the Future, Westlake University

## Abstract

Increasing works for antibody design are emerging to generate sequences and structures in Complementarity Determining Regions (CDRs), but problems still exist. We focus on two of them: *(i) authenticity of the generated structure* and *(ii) rationality of the affinity maturation*, and propose G_EO_AB as a solution. In specific, GeoAB-Designer generates CDR structures with realistic internal geometries, composed of a generative geometry initializer (Geo-Initializer) and a position refiner (Geo-Refiner); GeoAB-Optimizer achieves affinity maturation by accurately predicting both the mutation effects and structures of mutant antibodies with the same network architecture as Geo-Refiner. Experiments show that G_EO_AB achieves state-of-the-art performance in CDR co-design and mutation effect predictions, and fulfills the discussed tasks effectively.

## 1. Introduction

Antibodies are immune proteins that can bind to a kind of antigen protein so as to recognize and neutralize the pathogen (Basu et al., 2019). There are two heavy chains and two light chains in an antibody, leading the antibodies to Y-shaped proteins. In each chain, three variable regions determine the binding property of the antibody towards the antigen, which are called Complementarity Determining Regions (CDRs). Therefore, to design antibodies that bind to specific antigens with desirable properties, the key problem is to establish computational approaches to accurately identify the 3D structures and 1D sequence of the amino acids in the CDRs because the large combinatorial space of amino acid types results in infeasible and unaffordable validation of wet-lab experiments.

Recently, deep learning has revolutionized fields like drug discovery and protein design (Jumper et al., 2021; Dauparas et al., 2022; Lin et al., 2022; 2023). To be specific, increasing works are emerging for antibody design, which considers antigens and conserved regions (non-CDRs) as context information, to achieve the co-design of the sequence and structure of the target CDRs (Kong et al., 2023a; Jin et al., 2021). Besides, based on the co-design models, the optimization of the antibody by mutating amino acids in the CDRs to enhance the binding affinity can be realized, which is called affinity maturation (Cai et al., 2023). Iterative Target Augmentation (ITA) is proposed to fulfill the affinity maturation tasks (Yang et al., 2020), by iteratively adding co-design models’ prediction of the mutant antibodies structures and sequences with higher affinity predicted by another pretrained model to the training set and thus to guide and retrain the co-design models to generate antibodies of high affinity. However, several problems still exist, and we focus on two of them: *(i) authenticity of the generated structure* and *(ii) rationality of the affinity maturation*.

To illustrate problem *(i)*, we claim that in the structure design of the CDRs, these methods usually ignore geometry constraints of protein structures, such as peptide planar and inflexible geometries like bond lengths (Robinson et al., 2014), causing the unrealistic modeling of the CDR structures. Figure 1 gives an illustration of the distribution gap between partial inflexible internal geometries (bond length, angles, and inflexible torsions) and redundant ones (flexible torsions) of the structures generated by state-of-the-art methods and real-world proteins (Sec. 5.1 gives details).

**Figure 1:**
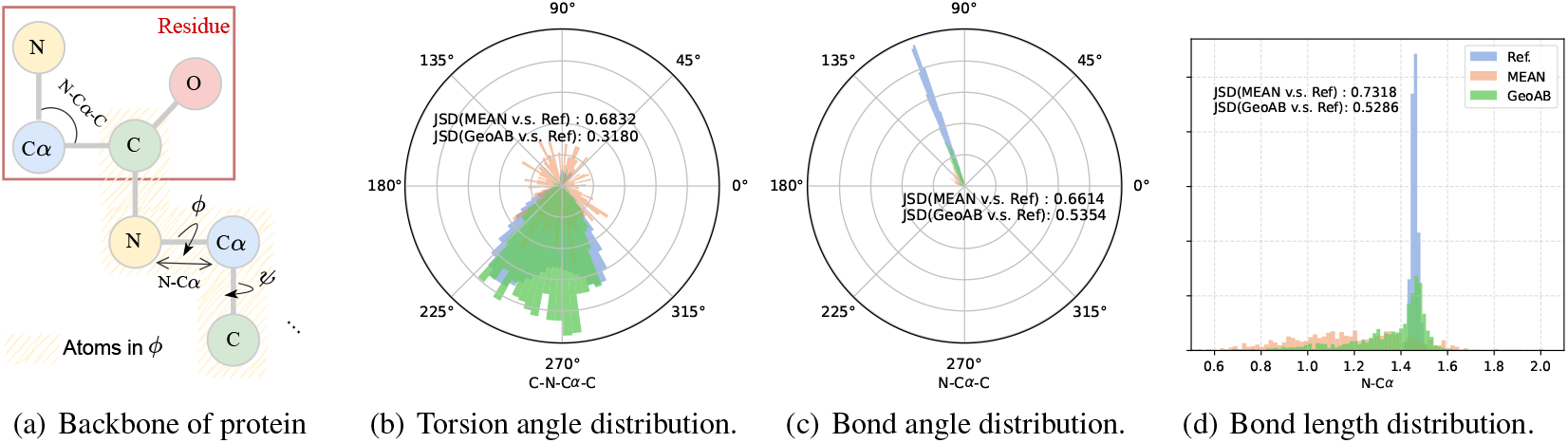
Distribution gap between several internal geometries of true protein structures as a reference and of generated ones. Jensen–Shannon divergence (JSD) is used as a measurement. (a) gives a sketch of the demonstrated geometries. (b,c,d) The torsion angle ‘C-N-C*α*-C’ is a redundant (flexible) geometry, while the bond angle ‘N-C*α*-C’ and length ‘N-C*α*’ are inflexible geometries. Internal geometries obtained from CDR structures generated by MEAN usually deviate from the true distribution greatly. In comparison, GeoAB narrows the gap and generates more realistic and reasonable CDR structures.

**Figure 2:**
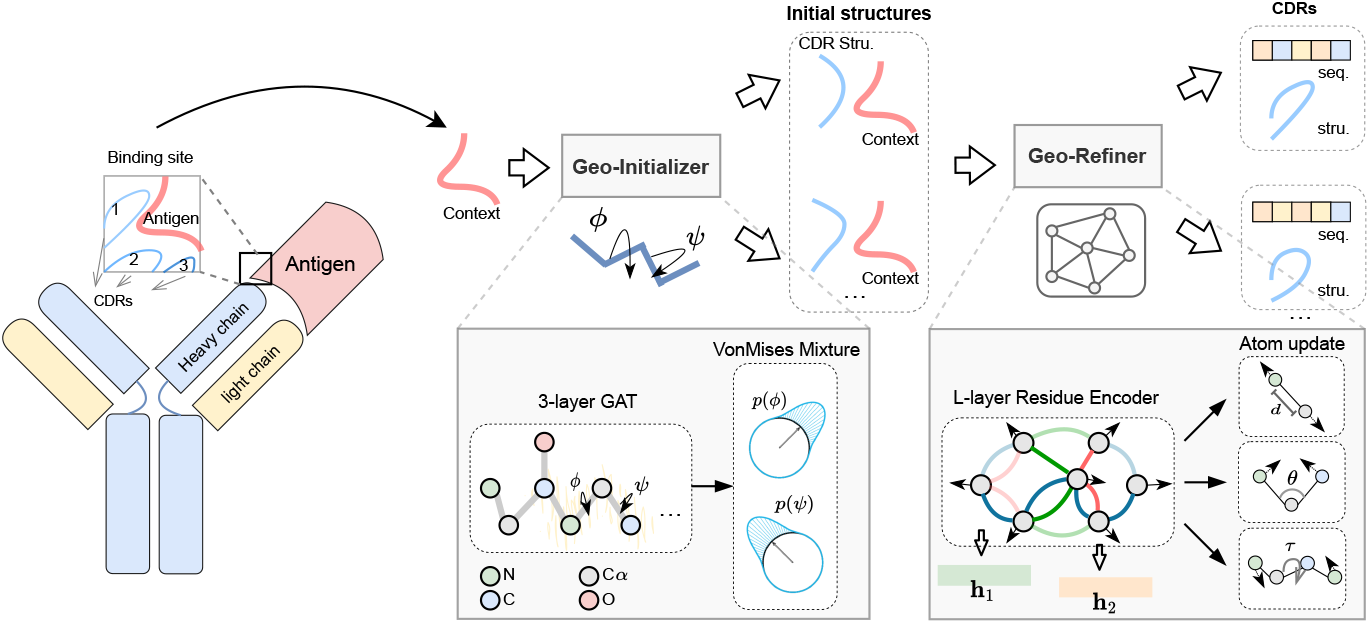
Workflows of **GeoAB-Designer** in antibody design task.

For problem *(ii)*, these methods first employ a trained model with sequences and structures as input to predict the change in binding free energy (ΔΔ*G*) and exploit the ITA algorithm to tackle the optimization. This procedure is irrational due to two problems. First, the used ΔΔ*G*-predictor requires mutant types’ structures as input, while they are usually unknown, so it uses predicted structures by methods like Rosetta programs (Park et al., 2016) as training instances. However, the generated mutant antibodies by the co-design models are of different domains from the training ones, resulting in the unreliability of the predicted ΔΔ*G* (See Fig. 5 as an example). Secondly, the inaccurate prediction of ΔΔ*G* cannot guide the ITA reliably to generate antibodies toward affinity maturation targets. The other approach for affinity maturation is to enumerate amino acid types on the points of interest. Previous works show that multiple-point mutations usually achieve successful affinity maturation (Sulea et al., 2018), but the enumeration of mutations on multiple points is unaffordable in wet-lab validation. Therefore, it is urgent to establish an effective computational method for narrowing down the search space for the task.

To address them, we proposed G_EO_AB. *On antibody CDR co-design*, it can jointly sample CDR sequences and structures with realistic protein geometry, by using heterogeneous residue-level encoder and equivariant atom-level interaction layers and employing energy-informed geometric constraints. In detail, GeoAB-Designer consists of a geometry initializer (Geo-Initializer) and a position refiner (Geo-Refiner). Geo-Initializer is a generative model with informative prior, which samples the redundant internal geometries and reconstructs the structures with NeRF (Parsons et al., 2005) . Further, Geo-Refiner as GNNs represents the binding site as hierarchical graphs, in which a highlevel graph models heterogeneous residue-level relations and a low-level one captures atoms’ interactions considering mechanics for bond lengths, bond angles, and torsion angles. Equivariance is ensured in updating the atom positions. *On mutation-based affinity maturation*, we propose a novel structure-aware GeoAB-Optimizer, based on the proposed network architecture and geometric constraints. It uses amino acid sequences and the context structures as input, and generates both structures of the CDRs and ΔΔ*G*, to avoid the problem of domain differences in input CDR structures. Based on the fact that the representations obtained by structure-related tasks usually assist energy-related prediction (Jin et al., 2023), we propose a structure-aware joint training strategy on paired labeled data (with ΔΔ*G* labels) and unpaired ones. Extensive experiments demonstrate the superiority of G_EO_AB in both realistic generation and reliable affinity maturation for antibody CDRs.

## 2. Background

### Notations

For a binding complex composed of an antigen-antibody pair as 𝒞, which contains *N*_aa_amino acids, there are *N*_ag_ amino acids in the antigen and *N*_cdr_ and *N*_ncdr_ amino acids of a particular CDR and other non-CDRs in the antibody. We represent their index set by ℐ_ag_, ℐ_cdr_ and ℐ_ncdr_ according to the sequential orders, where |ℐ_ab_| = *N*_ab_, |ℐ_cdr_ |= *N*_cdr_ and |ℐ_ncdr_| = *N*_ncdr_. For a protein, we consider four backbone atoms for each amino acid, so one amino acid can be represented by its type *a*_*i*_ ∈ {1, …, 20} and atom coordinates of{ N, C*α*, C, O} as (***x***_*i*,1_, …, ***x***_*i*,4_), where ***x***_*i,j*_ ∈ ℝ_3_. Therefore, 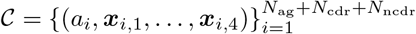 can be split into three sets as 𝒞 = 𝒢 ∪ ℛ ∪ 𝒩 according to the index sets of antigen, CDR and non-CDR. For each 𝒞, Gibbs free energy of association is the physical quantity used to measure the binding affinity between antigens and antibodies, denoted by Δ*G* = *E*(𝒞). When mutations occur, 𝒞= 𝒞 ^(wt)^ as wild types will be changed into 𝒞^(mt)^ as mutant types, with both structures and sequences changed. The mutation effect is usually measured by the change of Gibbs free energy as ΔΔ*G* = *E*(𝒞_(mt)_) *− E*(𝒞_(wt)_).

### CDR co-design

For CDR co-design, our goal is to establish a model to learn the distribution of CDRs conditioned on the antigens and non-CDRs of the antibodies, *i*.*e. p*(ℛ | 𝒢 ∪ 𝒩). For realistic structure design, the internal geometries of the protein structures generated by models should be close to the true ones that are governed by physicochemical rules. For example, the planarity of peptide bonds usually constrains the torsion angle of ‘O=C-N-C*α*’ to be 0, and steric collisions between atoms lead torsion angles of *ϕ* and *ψ* to fall into defined regions in a graph called the Ramachandran plot (Agnihotry et al., 2022) (Details in Appendix. A.1 ).

### Muation-based CDR affinity maturation

For affinity maturation, we aim to develop a model that can generate 𝒞^(mt)^ that satisfies *E*(𝒞 ^(mt)^) − *E*(𝒞 ^(wt)^) *<* 0, based on 𝒞^(wt)^ Since *E*(.) is intractable and the mutation data is limited, one key issue is the reliability of direct prediction of ΔΔ*G*(𝒞^(mt)^, 𝒞 ^(wt)^). Otherwise, the maturation is unreasonable given an inaccurate evaluation of the mutation effects. In addition, as a single mutation of amino acid types may affect the structures on a larger scale, it is also important to consider the structure flexibility of mutant types, since their structures are usually unknown. Note that CDRs make a disproportionately large contribution to the binding interactions in the antigen-antibody complexes, so we focus on the mutation effects of points located in the CDRs. Therefore, the problem can be formulated as to develop a model *q*(ℛ^(mt)^ 𝒞^(wt)^), *s*.*t*. ΔΔ*G*(𝒞^(mt)^, 𝒞 ^(wt)^) *<* 0 for ℛ ^(mt)^ ∼ *q*, based on our assumption that the structures of non-CDRs and antigens are inflexible.

## 3. Method

### 3.1 Graph Construction

We denote the antibody-antigen residue graph as heterogeneous graph (𝒱_aa_; ℰ_1_, ℰ_2_, ℰ_3_), where the nodes are denoted by the residues with C*α* coordinates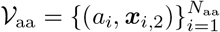, and the edge sets are composed of three kinds: sequence edge set ℰ_1_ obtained by whether the two amino acids are consecutive in the protein sequence, *i*.*e*. (*i, j*) ∈ ℰ_1_ if they are consecutive, intra- and inter-structure edge sets ℰ_2_ and ℰ_3_ obtained by whether the two amino acids are close in distance and belong to the same/different proteins. For atom graph (𝒱_at_; ℰ_at_), the nodes are represented by its atom types *e*_*j*_ *∈* {1, 2, 3, 4} and positions 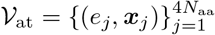, and the edge set ℰ_at_ is constructed according to the chemical bonds as Figure 1.(a) shows.

### 3.2 Geo-Refiner

Given the initial states of the backbone atom positions and amino acid types which are often inaccurate, the Geo-Refiner both refines the structures of the backbone atoms and predicts the amino acid types in CDRs. The refinement process is deterministic and includes two steps: residue-level encoding and atom-level updating.

### Heterogeneous residue-level encoding

In the residuelevel graph (𝒱_aa_; ℰ_1_, ℰ_2_, ℰ_3_), we use a simple heterogeneous GNN to encode the residues. Considering a *L*-layer heterogeneous residue encoder, the formulation of the *l*-th equivariant layer can be defined as follows:

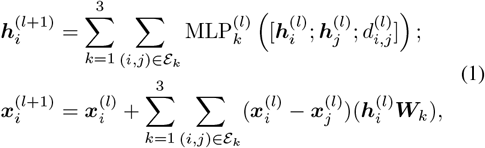

For 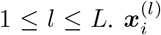 is *i*-th node’s updated position of C*α* atom after (*l−* 1) layers, and 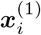 is the initialized C*α* positions which can be obtained by an initializer (see Sec. 3.3). 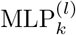 as a shallow multi-layer perceptron (MLP), encodes the *i* and *j* nodes’ embeddings and inter distance 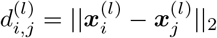 as messages, and summation is used to aggregate messages. ***W***_*k*_ projects embeddings of high dimension into one. 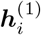 as the input of the GNN, is the initial embedding of residues, which considers amino acid’s sequence position *i*, types *a*_*i*_, intra-residues’ bond lengths, angles and torsion angles, and inter-residues’ distances, angles, and dihedrals.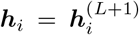 is the output E(3)invariant representation, which will be used for updating the atom representations and other downstream tasks.

### Mechanics-informed atom-level updating

In the atomlevel graph (𝒱_at_; ℰ_at_), atom representations are first encoded by ***y***_*i*_ = Emb(*e*_*i*_) + ***h***_*i*_, for 1 ≤*i≤*4*N*, where Emb(·) is a simple lookup table, and ***h***_*i*_ is the representation of the residue which the atom belongs to, in Eq. 1. A threelayer GAT (Veličković et al., 2018) is stacked for atom-level message passing, as 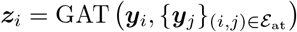.

In position updating, we generalize the graph mechanics networks (Huang et al., 2022) to binary, ternary, and quaternary interactions, and harness generalized coordinates to describe the kinematics of the atoms. In specific, for a group of connected atoms as 𝒮= {(***z***_*i*_, ***x***_*i*_)} _1≤*i*≤*M*_, position and velocity in the generalized coordinates as ***q***_*i*_ and 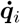, and velocity in global coordinates as 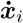, the updating process is

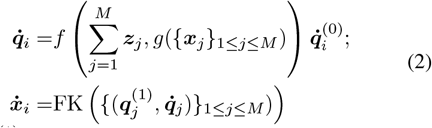

Where 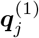 is the initial atom position in generalized coordinates, 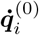 is velocity direction, *g*(. ) obtains the representation of invariant intra-geometries in the group, *f* : ℝ_3_ → ℝ is an arbitrary MLP, and forward kinematics FK(. ) projects the positions in updated generalized coordinates back to the global states. We detail how to update the position states in atom-level interaction layers for the three kinds.

To model the **binary interaction** with *M* = 2, we consider the two connected atoms indexed by (*i, j*). The two atoms connected by a chemical bond cannot form a generalized coordinate, and thus we use the global coordinate for updating the positions. The velocity direction 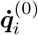 is parallel to the bond for scaling the bond length, as 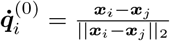, and 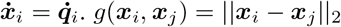 is the bond length. Note that (*i, j*) can be commutative, and the binary interaction is equivalent to vanilla EGNN (Satorras et al., 2021).

To model the **ternary interaction** with *M* = 3, we consider the three consecutively connected atoms indexed by (*i, j, k*). To update the bond angle *θ* = *θ*_*i,j,k*_, the center atom position ***x***_*j*_ is fixed, and 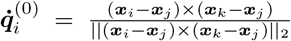 represents unit angular velocity. *g*(***x***_*i*_, ***x***_*j*_, ***x***_*k*_) = [sin (*θ*); cos (*θ*)] is the angle encoding. The forward kinematics written as 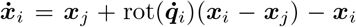 model the velocity in the global coordinates, where rot 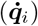 indicates the rotation matrix around the direction of 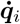 by absolute angle 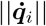 . The dynamic updates for atom *k* are similar.

To model the **quaternary interaction** with *M* = 4, the four consecutively connected atoms are indexed by (*i, j, k, l*). To update the torsion angle *τ* = *τ*_*i,j,k,l*_, the atom positions ***x***_*j*_ and ***x***_*k*_ are fixed, and the atoms *i* and *l* turn round on the axis (***x***_*j*_ *−* ***x***_*k*_), so that 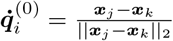 and 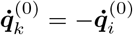 are unit torsion angular velocities. *g*(*·*) encodes torsion angles by [sin |*τ*|; cos |*τ*|], in which the absolute values of *τ* are taken to avoid parity (Jing et al., 2023). The FK(*·*) is written as 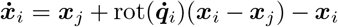 for atom *i* and 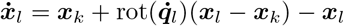 for atom *l*.

As each atom may belong to several groups, the final updates of the positions in the global coordinates are the summation of all the updates in different groups. Because the binary and quaternary interaction layers are both linear combinations of {***x***_*i*_} _1*≤i≤M*_, they are E(3)-equivariant (Villar et al., 2023); The cross product operation in ternary interaction layer leads it to be roto-translational but not reflectional equivariant (Geiger & Smidt, 2022), *i*.*e*. SE(3)-equivariant. In practice, we find Geo-Refiner does not need multi-round iterations, *i*.*e*. One-shot prediction achieves comparable performance.

### 3.3 Geo-Initializer

For antigens and non-CDRs, the structures of ground truth are used as initial states. For CDRs, linear interpolation is usually used to initialize atom positions (Kong et al., 2023a), where the equispaced atoms are distributed according to the (min ℐ_cdr_ *−* 1)-th and (max ℐ_cdr_ + 1)-th amino acids’ positions. However, the initialization cannot give any informative prior knowledge for realistic structure generation.

To initialize realistic structures of CDRs, we propose Geo-Initializer to directly generate internal geometries. We observed that the redundant geometries in modeling the backbone atoms are three torsion angles of ‘N-C*α*-C-N’, ‘C*α*-CN-C*α*’ and ‘C-N-C*α*-C’ (Appendix. A.2). Therefore, we use Von Mises distributions to generate the three torsion angles (Swanson et al., 2023), since it is a continuous probability distribution on the circle with support in [*− π, π*). As the multi-modal nature of rotatable bond torsion angles in ‘NC*α*-C-N’ and ‘C-N-C*α*-C’ is observed (Appendix. B.1), a mixture of *K* von Mises distributions is employed to capture the modality, as 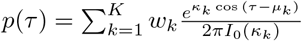, in which *τ* is the torsion angle, *I*_0_(*·*) is the modified Bessel function of order 0, *w*_*k*_ is the weight, *μ*_*k*_ is the mean, and *κ*_*k*_ is the concentration of the *i*-th distribution. To obtain the parameters of 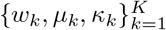, a shallow GAT is used to encode the atom-level graph to obtain representation for each atom. After summing the representations of the four atoms that make up the torsion angle *τ*, three shallow MLPs project the summation to *K*-dimensions, leading to 3 *× K* predicted parameters in *p*(*τ* ). To train the Geo-Initializer, we minimize the negative log-likelihood of the ground truth torsion angle samples for a given rotatable bond under a mixture of *K* Von Mises distributions defined by the predicted parameters.

Geo-Initializer thus is a generative model from which the redundant torsions of the backbones are sampled. For the rest of the inflexible geometries, the ideal values are used. In this way, NeRF is employed to reconstruct the backbone atom positions as the initial structures for further refinements.

### 3.4 Geometric Constraints

For structure refinement, the outputs of atom-level interaction layers are the backbone atom positions in CDRs in global coordinates, denoted by *𝒳*, and the superscripts ^0^ denote the true values to differentiate the predictions. Inspired by AMBER (Maier et al., 2015), we propose to incorporate the following energy terms as geometry constraints, to govern the generated structure with chemical rules, including constraints on bond length, angles, and torsion angles.

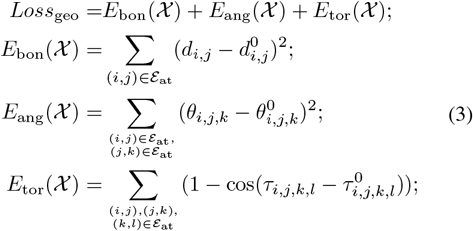

where *d*_*i,j*_ is the bond length between 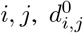 denotes the expected bond length between atom *i, j*, which can be obtained by the true structures. For *θ*_*i,j,k*_ and 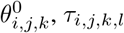and 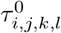, the definitions are similar.

Besides, two loss functions are employed: (i) residual-level error on C*α* positions and (ii) atom-level error on all backbone atoms. Since the objective region for designing is CDRs, for *i* ∈ ℐ_cdr_, the loss for position can be written as

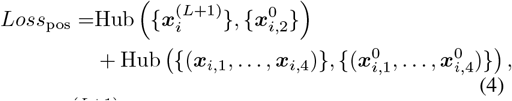

Where 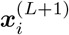 is the predicted position of residue *i*, output by *L*-layer residue-level encoders and defined as C*α*’s positions in Eq.1, and 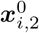 is the true C*α*’s position of residue *i*; (***x***_*i*,1_, …, ***x***_*i*,4_) is the four backbone atoms of residue *i*, obtained by atom-level interaction layers, and 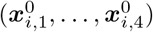 is that of ground truth. Hub(*·* ) is the Huber loss used for training stability. The designated position loss aims first to determine the coarse positions of residues and then refine the grained positions of the backbone atoms.

### 3.5 Antibody CDR Co-Design

In the task of antibody CDR design, we aim to co-design both the amino acid sequence and the backbone atom’s structures in CDRs. For sequence design, a masked state as an absorbing type is used to initialize the amino acid types. For structure design, we propose two schemes for the task: (i) **GeoAB-Designer** composed of ‘(pre-trained) Geo-Initializer + Geo-Refiner’ as a generative model due to the stochasticity of Geo-Initializer and (ii) **GeoAB-Refiner** composed of ‘Linear Initialization + Geo-Refiner’ as a refinement model. By this means, the loss function is

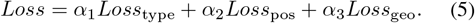

{*α*_*j*_}_*j*=1,2,3_ are loss weights. 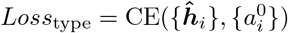, in which 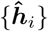 are the logits, obtained by a linear classifier stacked after the *L*-layer residue encoder, projecting 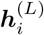 from *D*-dimension to 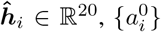 are the true amino acid types in CDRs, and CE(*·*) is the cross entropy loss.

### Dynamic weights

In **GeoAB-Designer**, the inflexible internal geometries in the initialized backbone structures are almost identical to the real ones (see Appendix. C.3). To preserve the initialized realisticity of the geometries, we hope that in the training stage, the constraints on internal geometries as *Loss*_geo_ can be more emphasized at the beginning, and gradually decreasing losses in positions *Loss*_pos_ in the training process can also lead to a further reduction in *Loss*_geo_, given that when the model can make perfect predictions of position such that *Loss*_pos_ *→* 0, *Loss*_geo_ is also minimized. Motivated by this, we use a training trick of dynamic loss weights, by setting *α*_3_ large at the beginning and gradually making it smaller with exponential moving average (See Appendix. B.2), to keep the geometry from deviating too much from the initial state during training.

### 3.6 Mutation-Based Affinity Maturation

For affinity maturation, two sequences of amino acids are given, *the wild type* and *the mutant type*. The reliable maturation requires our model to accurately predict both the structures and the mutation effect ΔΔ*G*. To tackle the problem, we propose an architecture composed of twin Geo-Refiners with shared parameters and a joint training strategy to make full use of the paired labeled and unpaired data.

In detail, we first assume that the mutant amino acid will affect the surrounding structures around it, but keep the rest unchanged, so we aim to predict the structure of the surrounding *N*_flx_ residues. A simple ‘Linear Initialization’ initializes the backbone atoms in the affected regions. For an instance of *paired data* with two types of sequence, the initial amino acid sequences as inputs are the true lists of types, and we write the index set of mutant points as ℐ_mt_. Two Geo-Refiners with shared parameters will generate the backbone structure of two types in the affected regions by the atom-level interaction layers. Besides, following (Shan et al., 2022), ΔΔ*G* is predicted by

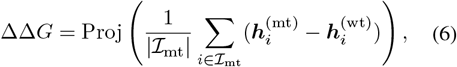

in which Proj(*·* ) is an MLP, projecting *D*-dimensional representation to a scalar. In this way, the loss function reads

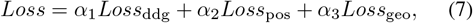

Where *Loss*_ddg_ = MSE(ΔΔ*G*,ΔΔ*G*^(0)^), and ΔΔG^(0)^ is the labeled change in Gibbs free energy. *Loss*_pos_ and *Loss*_geo_ are usually calculated with the structures of wild types since the structures of the mutant ones are usually unavailable. For an instance of *unpaired data* with a single type of sequence that is usually defined as wild type, we select a point of interest and mask its surrounding *N*_flx_ residuals structure by the same initialization method. The mutant type is defined as itself, and the loss on mutation effect will change into *Loss*_ddg_ = MSE(ΔΔ*G*, 0).

Once the model is trained, we can obtain the mutant sequences and structures such that ΔΔ*G <* 0 by enumerating the other 19 amino acids and replacing the point of interest.

## 4. Related Work

### Antibody Design

Classical works are usually based on hand-crafted potential functions, including physical forcefield terms (Li et al., 2014; Lapidoth et al., 2015; Adolf-Bryfogle et al., 2018) and statistical potential (Min-yi & Andrej, 2006). This results in intensive computation and unavoidable inaccuracy, as complex mechanisms of protein structure cannot be described by simple potentials. Deeplearning-based methods concentrated on 1D sequence design (Alley et al., 2019; Liu et al., 2020; Saka et al., 2021; Akbar et al., 2022), and then, thanks to great process in graph neural networks (Wu et al., 2021; Liu et al., 2021; Wu et al., 2023) and protein modeling (Huang et al., 2023; Wu et al., 2024; Tan et al., 2024; 2023), more works have been focused on structure-sequence co-design. For example, RefineGNN (Jin et al., 2021) auto-regressively generate the amino-acid types and structures; HERN (Jin et al., 2022) achieves co-design and docking tasks by using a hierarchical graph; MEAN (Kong et al., 2023a) employ multi-channel equivariant attention networks to generate CDRs with multiround refinements; DyMEAN (Kong et al., 2023b) similarly can fulfill the co-design tasks, and simultaneously dock to the epitope at a full-atom level. DiffAB as diffusion models generates translation, orientation, and type variables (Luo et al., 2022). HTP (Wu & Li, 2023) and ABGNN (Gao et al., 2023) focus on antibody pretraining. These methods hardly generate antibodies that conform to physicochemical constraints, while GeoAB focuses on realistic generation.

### Mutation Effect Prediction

Similarly, traditional approaches utilize energy functions to model the interactions and compute the ΔΔ*G* (Steinegger & Söding, 2017). Statistical methods rely on feature engineering and use the invariant features and statistical learning methods to predict mutation effects (Lei et al., 2023). In deep learning, the effectiveness in predicting ΔΔ*G* is validated. For example, ESM-1v (Meier et al., 2021) proposes sequence-based pretraining tasks and predicts mutation effects on protein functions. DDG-Predictor (Shan et al., 2022) takes structures and sequences of both types as input, and predicts the ΔΔ*G* effectively. However, the structures of mutant types are usually unknown, RDE-PPI (Luo et al., 2023) avoids the problem and uses a structure-aware pretrained network on side-chain rotamers to assist the prediction. Besides, there are also works on designing pretraining tasks for ΔΔ*G* predictions (Hsu et al., 2022; Yang et al., 2023; Cai et al., 2023). In comparison, G_EO_AB also considers mutation effects attributing to structural flexibility but uses joint training strategies instead of pretraining paradigms.

## 5. Experiment

### 5.1 Antibody CDR Co-Design

#### Baselines

We select 6 methods as benchmarks. For refinement methods, **RefineGNN, MEAN** and **DyMEAN** are used. Their variants of **C-RefineGNN** and **C-DyMEAN** are included, meaning that the two methods use the same contextual condition. RefineGNN only models the heavy chains with less context and DyMEAN is designed for both docking and co-designing with the unknown paratope. For generative methods, we choose **DiffAB** as a baseline, which generates translation, orientation, and type variables, with the intra-residue geometries fixed with ideal values. As discussed in Sec. 3.5, we propose two variants of GeoAB. The first is **GeoAB-Refiner** in comparison with the refinement methods; the second is **GeoAB-Designer**, as a generative model compared with DiffAB. For GeoAB, our model is open to the public through https://github.com/Edapinenut/GeoAB.

#### Setup

Following (Kong et al., 2023a), we use the SAbDab (Dunbar et al., 2014) with complete antibody-antigen structures for training. IMGT scheme is used to renumber the complexes (Lefranc et al., 2003). The splits of training, validation and test sets are according to the clustering of CDRs via MMSeqs2 (Steinegger & Söding, 2017). After the 10 cross-validations on SAbDab, we benchmark all methods with the 60 diverse complexes as RAbD (Adolf-Bryfogle et al., 2018), and give comparisons. The training is still conducted on the SAbDab dataset, but we eliminate all antibodies from SAbDab whose CDR-H3s share the same cluster as those in RAbD to avoid potential data leakage. We use AAR metrics to evaluate amino-acid recovery ratio; A-RMSD and UA-RMSD as the RMSD between C*α* with/without Kabsch alignment; lddt and TM-score measuring global structural similarities. For the generative methods, group comparisons are made by selecting percentile of 20%, 50%, and 100% according to UA-RMSD. For details see Appendix. C.1.

#### Results

Table. 1 gives results on RAbD CDR-H3 designing, and for 10-fold cross-validation on SabDab, it is given in Appendix. C.2. It shows that in refinement models, GeoAB-R achieves the overall best performance, especially in structure evaluation, with 10.71% and 10.02% improvements in A-RMSD and UA-RMSD respectively over state-of-the-art C-DyMEAN. In generative models, GeoAB-D also outperforms DiffAB in all aspects by a large margin.

**Table 1:**
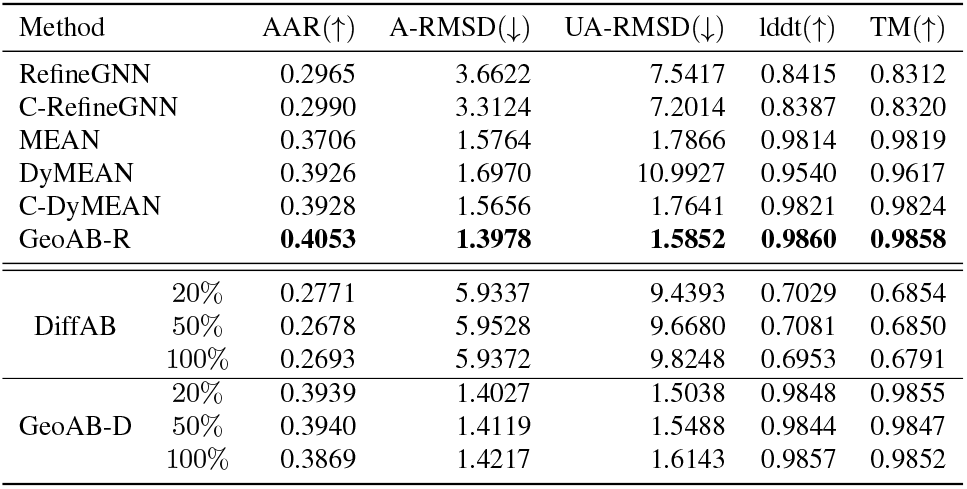
Metrics on generated CDR-H3 for RAbD compared with the reference. ‘GeoAB-R’ is the refinement variant and ‘GeoAB-D’ is the generative one. Values in **bold** are the best metrics.

#### Geometry analysis

Since GeoAB aims to generate realistic structures, we analyze internal geometries. We use Jensen–Shannon divergence (JSD) to evaluate the distribution differences and use mean absolute error (MAE) and mean cosine error (MCE) as two metrics to measure the instance differences in lengths and angles, respectively. In specific, MCE is written as MCE(*θ*^0^, *θ*) = 1*−* cos(*θ*^0^*− θ*), reaching 0 when *θ* = *θ*^0^( mod 2*π*). Table. 2 gives the comparison, including inflexible geometries like bond lengths ‘C*α*-C’ and ‘C-N’, bond angles ‘N-C*α*-C’ and ‘C*α*-C-N’, and torsion angles ‘O=C-N-C*α*’ and flexible torsions ‘CN-C*α*-C’. GeoAB-D generates the most realistic internal geometries in overall evaluations. For inflexible geometries, because DiffAB regards the residues as rigid bodies, the intra-residue geometries are set as ideal values, leading to the least deviation from references in ‘C*α*-C’ and ‘N-C*α*-C’. However, in inter-residue geometries such as ‘C-N’ and ‘CN-C*α*-C’, GeoAB shows superiority over DiffAB. Further, GeoAB-D as a generative model, initialized by the prior torsion angle initializer, outperforms GeoAB-R in JSD, but is less competitive in MAE/MCE due to its stochasticity. Figure. 1 shows the distribution gap between GeoAB-R and MEAN; Figure. 4 gives an example to show that GeoAB-D generates more realistic CDR-H3 structures than MEAN.

**Table 2:** Metrics on flexible and inflexible geometries obtained by generated CDR-H3 *v*.*s*. reference. Values in **bold** are the best metrics, and in underline are the second. The full comparison is given in Appendix. C.3.

| Geometry | JSD(↓) | | | | | MAE/MCE(↓) | | | | |
| --- | --- | --- | --- | --- | --- | --- | --- | --- | --- | --- |
|  | MEAN | C-DyMEAN | DiffAB | GeoAB-R | GeoAB-D | MEAN | C-DyMEAN | DiffAB | GeoAB-R | GeoAB-D |
| Cα-C | 0.7318 | 0.7673 | **0.1582** | 0.5286 | <u>0.5084</u> | 0.3209 | 0.5310 | **0.0298** | 0.2182 | <u>0.1883</u> |
| C-N | 0.6720 | 0.7350 | 0.6537 | <u>0.5916</u> | **0.5477** | 0.3824 | 0.4784 | 0.6202 | <u>0.3201</u> | **0.2537** |
| N-Cα-C | 0.6614 | 0.5646 | **0.3207** | 0.5354 | <u>0.5309</u> | 0.2512 | <u>0.1646</u> | **0.1239** | 0.1853 | 0.2207 |
| Cα-C-N | 0.6819 | 0.7583 | 0.6108 | <u>0.6031</u> | **0.5628** | 0.2368 | 0.5441 | 0.2181 | **0.1479** | <u>0.1908</u> |
| O=C-N-Cα | 0.6015 | 0.6670 | 0.4354 | <u>0.4193</u> | **0.3657** | 0.7799 | 0.9813 | 0.8524 | <u>0.3158</u> | **0.2992** |
| C-N-Cα-C | 0.6832 | 0.3506 | 0.4992 | **0.3180** | <u>0.3250</u> | 0.5611 | 0.9952 | 0.7277 | **0.3313** | <u>0.3634</u> |

### 5.2 Mutation-Based Remodeling

#### Baselines

We compare our method with 4 basedline ΔΔ*G*predictors, including **Rosetta** ddG (Alford et al., 2017; Leman et al., 2020), **FoldX** (Delgado et al., 2019), **DDG-Pred** and **RDE-PPI**. The first two are classical energy-based, and the last two are structure-based deep models. Note that all these methods are not pretrained, and **RDE-PPI** is tested as an unpretrained version for fair comparisons.

#### Performance Evaluation

Following (Luo et al., 2023), we use SKEMPI2 (Jankauskaite? et al., 2019) as the evaluation datasets. The datasets are split into three folds by structure, in which two of them are used for training and validation, and the rest are used for testing. Cross-validation is evaluated for ΔΔ*G*-predictors. Four metrics including Pearson correlation coefficient (PCC), Spearman’s rank correlation coefficient (SRCC), RMSE, and MAE are used to measure the accuracy of predictions. Besides, the per-structure PCC and SRCC are calculated in each structure of the protein complex. Table. 3 shows that GeoAB achieves competitive performance in the ΔΔ*G* prediction tasks. In addition, GeoAB does not require the structures surrounding the mutant points as input, and predicts both the structures and ΔΔ*G*, so the domain difference problems resulting from the unavailability of mutant types’ structures are avoided.

**Table 3:**
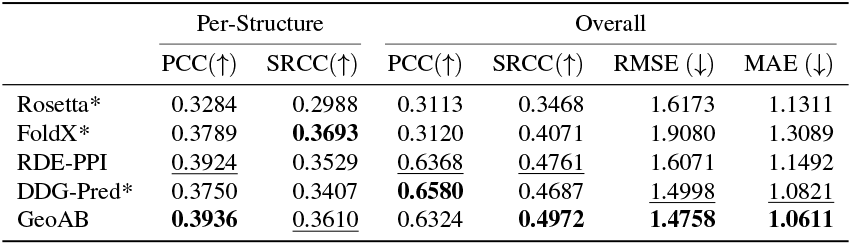
Metrics for cross-validation evaluation of ΔΔ*G* prediction on SKEMEPI2 dataset. Results of the methods with ‘*’ are from RDE-PPI (Luo et al., 2023). Values in **bold** are the best metrics, and in underline are the second.

|  | Per-Structure |  | Overall |  |  |  |
| --- | --- | --- | --- | --- | --- | --- |
| | PCC(↑) | SRCC(↑) | PCC(↑) | SRCC(↑) | RMSE (↓) | MAE (↓) |
| Rosetta* | 0.3284 | 0.2988 | 0.3113 | 0.3468 | 1.6173 | 1.1311 |
| FoldX* | 0.3789 | **0.3693** | 0.3120 | 0.4071 | 1.9080 | 1.3089 |
| RDE-PPI | <u>0.3924</u> | 0.3529 | <u>0.6368</u> | <u>0.4761</u> | 1.6071 | 1.1492 |
| DDG-Pred* | 0.3750 | 0.3407 | **0.6580** | 0.4687 | <u>1.4998</u> | <u>1.0821</u> |
| GeoAB | **0.3936** | <u>0.3610</u> | 0.6324 | **0.4972** | **1.4758** | **1.0611** |

#### Affinity Maturation

After ensuring our models’ ability of reliably predicting ΔΔ*G*, we retrained it with the joint training strategy. In detail, we divide the SKEMPI2 datasets into two sets, where the test instances are the antibody-antigen complexes that appear in the RAbD, and the rest of SKEMPI2 as *paired data* together with the training instances in SAbDab as *unpaired data* make up of the training set. PDB-REDO (Joosten et al., 2014) are also used as *unpaired data* included in the training set. For training instance in SKEMPI2, we set *N*_flx_ *∼ U* (11, 15) which mimics the length of CDR-H3 regions and masks the structures of the *N*_flx_ residues surrounding the mutant points by linear interpolation initialization, and Geo-Refiner is used to predict the masked structures and ΔΔ*G* as discussed in Sec. 3.6. For the training instance in SAbDab, the masked region is CDRH3, and the mutant types are regarded the same as the wild types, leading to ΔΔ*G* = 0. For PDB-REDO, we randomly select one residue on a chain and mask the surrounding *N*_flx_ residues’ structures for prediction, utilizing more data to enable the model to perceive the local structures.

#### Maturation Results

Table. 4 gives performance on the accuracy of predicting ΔΔ*G* and structures on the test set on SKEMPI2. For affinity maturation tasks, we conduct the optimizing processes as shown in Figure. 3, by randomly choosing 1 or 2 points in the CDR-H3 regions and enumerating the sequences as the model inputs 400 times for each antibody-antigen complex. The sequences and the predicted structures with the lowest ΔΔ*G* predictions are selected as the optimized samples with affinity maturation. We also give the average ΔΔ*G* of the maturation results on the 60 complexes in Table. 4, with results of ITA with GeoAB-Refiner also provided. Figure. 5 gives a sample optimized by **GeoAB-Optimizer**, and an unrealistic sample optimized by MEAN with ITA, where ΔΔ*G* is predicted by DDGPred in ITA. It shows that the training set of ΔΔ*G*-predictor (generated by Rosetta) is of great difference from the testing instance (generated by MEAN), so it gives a low ΔΔ*G* even if the generated structure is very unrealistic (broken chains in CDRs). In comparison, our **GeoAB-Optimizer** avoids the problem because it does not need the mutant type’s structures as input. Besides, the detailed results of **ITA** are given in Appendix. C.4 for fair comparisons.

**Table 4:** The results for struture and ΔΔ*G* prediction o SKEMEPI2 test set and affinit maturation on RAbD CDR-H3.

|  |  | SKEMPI2 | RAbD |
| --- | --- | --- | --- |
|  | A-RMSD | 3.4733 | 1.5982 |
|  | UA-RMSD | 2.6785 | 1.4006 |
| Per-Stru | PCC | 0.3977 | / |
|  | SRCC | 0.3845 | / |
| Overall | PCC | 0.6657 | / |
|  | SRCC | 0.5122 | / |
| Optimized $\Delta\Delta G$ | | / | -3.1598 |
| Optimized $\Delta\Delta G$ (ITA) | | / | -7.2816 |

**Figure 3:**
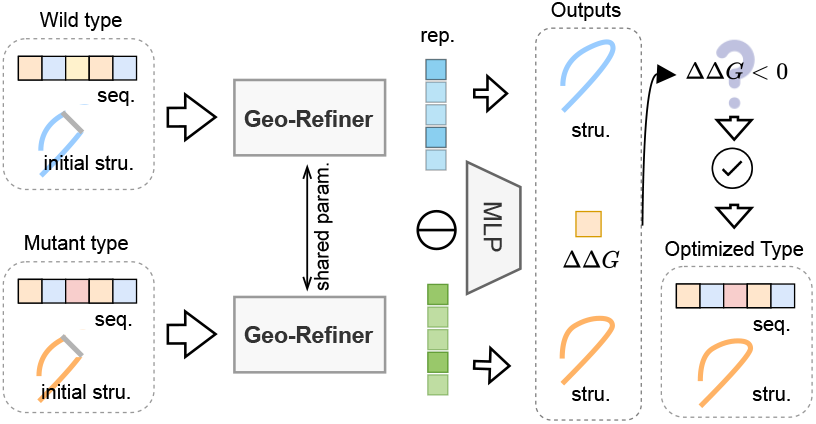
Workflows of **GeoAB-Optimizer** in antibody affinity muturation task.

**Figure 4:**
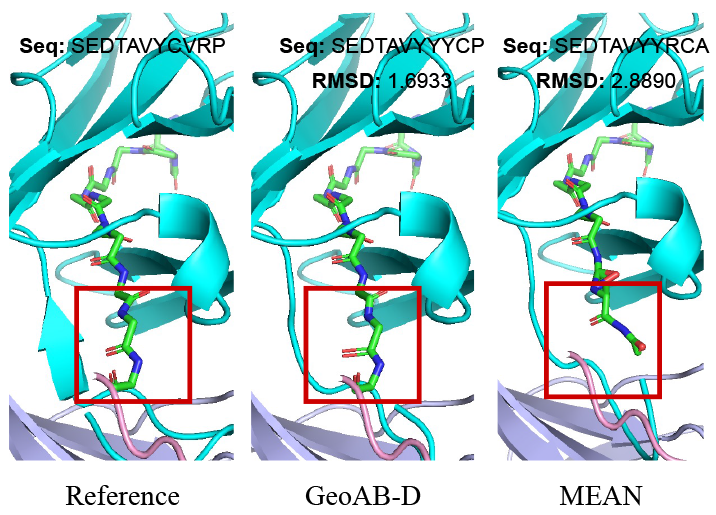
Backbone atoms in CDR-H3 (shown in sticks) on PDB-`3nid`, of the reference and designed by GeoAB and MEAN. RMSDs are calculated with all backbone atoms.

**Figure 5:**
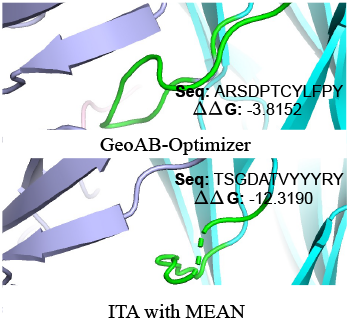
Optimized CDR-H3s for PDB-`4u6h`.

## 6. Analysis

We test if each proposed technique is necessary in GeoAB. We conduct ablation studies on SAbDab training and RAbD testing following the *setup in Table. 1*. In Table. 5, ‘w/o Bin/Ter/QuaIL’ means GeoAB-R without Binary or Ternary or Quaternary interaction layers for position update; ‘w/o Bon/Ang/TorGC’ means GeoAB-R without Bond length or Angle or Torsion geometric constraints; ‘w/o DyLW’ means training GeoAB with or without dynamic loss weight tricks; Besides, we also conduct experiments on the effects of ESM pretraining embedding, as ‘w/o ESM’. We choose representative metrics including AAR, UA-RMSD, and JSD in ‘C*α*-C’, ‘C*α*-C-N’, and ‘C-N-C*α*-C’ to illustrate how these techniques affect the designation. It can be concluded from Table. 5 that (1) The interaction layers help to improve UA-RMSE through a better prediction on flexible torsion angle of ‘C-N-C*α*-C’ except that ‘TerIL’ contributes little ; (2) Realistic inflexible geometries are usually generated by models with corresponding geometric constraints, according to ‘w/o BonGC’ and ‘w/o AngGC’. ‘TorGC’ also benefits the torsion angle prediction; (3) ESM embedding brings little improvements to the co-design tasks; (4) ‘DyLW’ helps GeoAB-G to generate more realistic inflexible geometries, since the JSDs are close to GeoAB-R when it is removed.

**Table 5:**
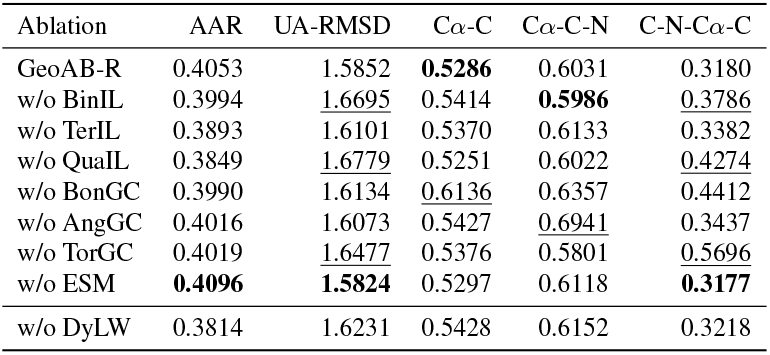
Ablation on proposed techniques. Geometries like ‘C*α*-C’ are JSDs of them. Values in **bold** are the best, and values in underline deviate from GeoAB-R obviously.

| Ablation | AAR | UA-RMSD | Cα-C | Cα-C-N | C-N-Cα-C |
| --- | --- | --- | --- | --- | --- |
| GeoAB-R | 0.4053 | 1.5852 | **0.5286** | 0.6031 | 0.3180 |
| w/o BinIL | 0.3994 | <u>1.6695</u> | 0.5414 | **0.5986** | <u>0.3786</u> |
| w/o TerIL | 0.3893 | 1.6101 | 0.5370 | 0.6133 | 0.3382 |
| w/o QuaIL | 0.3849 | <u>1.6779</u> | 0.5251 | 0.6022 | <u>0.4274</u> |
| w/o BonGC | 0.3990 | 1.6134 | <u>0.6136</u> | 0.6357 | 0.4412 |
| w/o AngGC | 0.4016 | 1.6073 | 0.5427 | <u>0.6941</u> | 0.3437 |
| w/o TorGC | 0.4019 | <u>1.6477</u> | 0.5376 | 0.5801 | <u>0.5696</u> |
| w/o ESM | **0.4096** | **1.5824** | 0.5297 | 0.6118 | **0.3177** |
| w/o DyLW | 0.3814 | 1.6231 | 0.5428 | 0.6152 | 0.3218 |

## 7. Concusion and Limitation

A method called GeoAB is proposed, which focuses on realistic CDR structure design and reliable affinity maturation. Two models are established as solutions. GeoAB-Designer is able to generate realistic CDR structures; GeoAB-Optimizer used for affinity maturation is able to predict ΔΔ*G* accurately and generate CDR structures, avoiding the problem of domain differences in input structures.

Still, limitations exist. First, the proposed interaction heads require more computational complexity than the vanilla EGNN, for each requires extra 𝒪 (*m*) updates on atoms’ positions, for a CDR composed of *m* corresponding groups. Secondly, the accuracy of prediction ΔΔ*G* is not as good as the models with structure-aware pretraining paradigms like RED-PPI, which will be our future concentration.

## Ackownledgements

This work was supported by the Science & Technology Innovation 2030 Major Program Project No. 2021ZD0150100, National Natural Science Foundation of China Project No. U21A20427, Project No. WU2022A009 from the Center of Synthetic Biology and Integrated Bioengineering of Westlake University, and Project No. WU2023C019 from the Westlake University Industries of the Future Research. Finally, we thank the Westlake University HPC Center for providing part of the computational resources. Besides, we thank the help of Dr. Tailin Wu, and his great efforts in his deep insights into cutting-edge issues, the guidance provided in the rebuttal, and the funding that supplied us with the necessary equipment for our research.

## Impact Statement

The world is revolutionizing itself in one epidemic after another. With the outbreak and long-term continuation of COVID-19, more and more AI scientists are beginning to turn their research interests to drug design. The focus of this paper is on one of them – antibody drugs. There is some work aimed at using AI algorithms to design and optimize antibodies. However, antibodies are a type of protein and few work focuses on the quality of their generation. Here, the present work presents a considerable contribution to improving both the quality of antibody generation and the reliability of optimization. We emphasize the social significance of the work here and hope to get the attention of the reviewers and the review committee.

## A. Protein Backbone Geometry Analysis

### A.1 Flexible and Inflexible Geometries

The topology of a part of proteins consists of residues, which are shown in Figure. 6(a) (Lieberman, 2014). The bond lengths and bond angles are ideally inflexible and governed by physicochemical laws, with the ideal values shown in Figure. 6(a). However, in molecules, the torsion angles are usually flexible, leading to the conformation flexibility of biomolecules. In the protein backbones, there are four main torsion angles, including *ϕ* as ‘C-N-C*α*-C’, *ψ* as ‘N-C*α*-C-N’, *ω* as ‘C*α*-C-N-C*α*’, and ‘C-N-C*α*=O’. Besides, ‘C-N’ as the peptide bond, usually has two states: trans, *ω ≈* 180^*°*^, and cis, *ω≈* 0^*°*^. In the trans configuration, the two alpha carbon atoms of the connected amino acids are on the opposite sides of the peptide bond, whereas in cis configuration they are on the same side of the peptide bond. In most cases, the peptide bonds in proteins are trans. This preference can be explained by the steric clashes that occur between groups attached to the alpha carbon atoms in cis form, which hinder the formation of this configuration. G.N. Ramachandran recognized that steric collisions between atoms prevent some combination of *ϕ* and *ψ* angles and, for the trans configuration, ranges of *ϕ* and *ψ* angles fall into defined regions in a graph called the Ramachandran plot as Figure. 6(b) shows. According to permitted *ϕ ψ* and *ω* angles, preferred conformations of the main chain lead to recurrent structures in proteins namely alpha helix, beta sheets, and turns (Cacabelos et al., 2014). Besides, due to the peptide planar, the ‘C*α*-C-N-C*α*’ torsion angles are also restricted as 0^*°*^.

**Figure 6:**
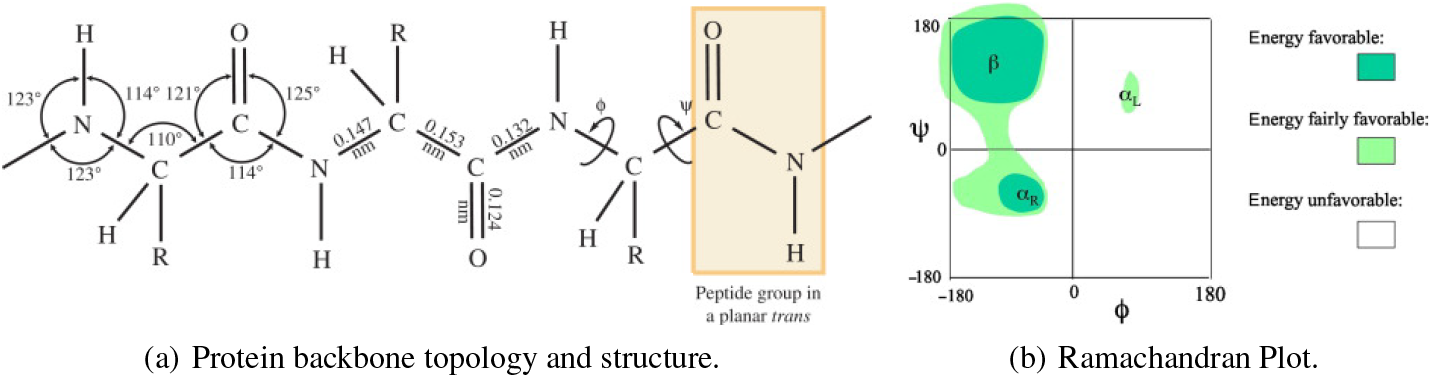
Illustration of peptide planar and the redundant geometry.

In this way, we can conclude that the flexibility of the backbone is mainly determined by the fluctuations in torsion angles, specifically *ϕ* and *ψ*, and the four flexible torsion angles degenerate into two. However, in real-world observation, *ω* are not strictly 0^*°*^ or 180^*°*^, generating a deviation of ±7^*°*^ from ideal values, which is also redundant geometry in accurate backbone generation tasks (MacArthur & Thornton, 1996). By this means, to generate a backbone, one can establish a generative model on the three torsion angles, such as FoldingDiff (Wu et al., 2022). Together, with the ideal values of inflexible geometries and generated redundant torsions, the protein backbone structures can be reconstructed by NeRF (Parsons et al., 2005), by sequentially placing the atom in the local coordinate systems, where the positions are determined by the length as distance, angles and dihedrals in the spherical coordinate systems.

### A.2 Analysis on SAbDab CDRs

Beyond the general proteins, we now focus on the antibodies’ CDRs structures, to figure out if the highly flexible regions follow the same rules as the former conclusions.

We first intercept the CDRs of all antibodies in the training set, and obtain empirical and kernel density estimated distributions of the chemical bond lengths, bond angles, and torsion angles of individual fragments, in Figure. 7.

**Figure 7:**
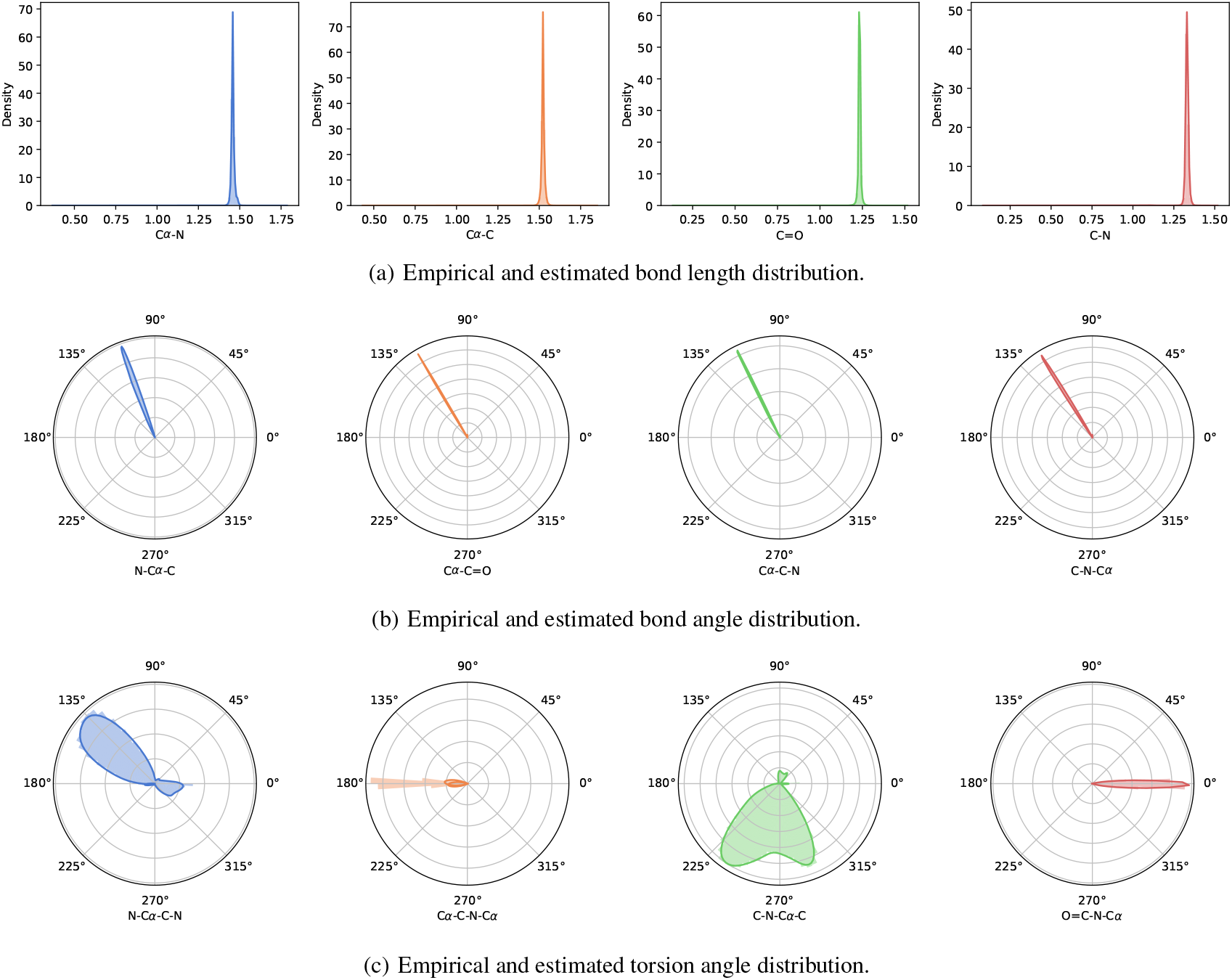
Distributions of flexibile and inflexible geometries obtained by CDRs in SabDab datasets.

It shows that the internal geometries basically follow the conclusions obtained in the last section. Distributions of *ϕ, ψ* and *ω* show the highest stds among all the 12 geometries, and all the *ω* torsion is trans. Besides, we give the statistical values on MAE between the ideal values and the observed ones and stds of each distribution, to further prove the conclusions in Table. 6. The same conclusion can be reached.

**Table 6:**
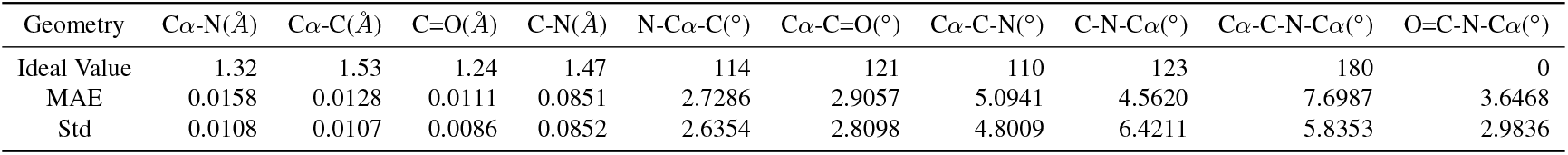
Statistics of flexible and inflexible geometries.

| Geometry | Cα-N(Å) | Cα-C(Å) | C=O(Å) | C-N(Å) | N-Cα-C(°) | Cα-C=O(°) | Cα-C-N(°) | C-N-Cα(°) | Cα-C-N-Cα(°) | O=C-N-Cα(°) |
| --- | --- | --- | --- | --- | --- | --- | --- | --- | --- | --- |
| Ideal Value | 1.32 | 1.53 | 1.24 | 1.47 | 114 | 121 | 110 | 123 | 180 | 0 |
| MAE | 0.0158 | 0.0128 | 0.0111 | 0.0851 | 2.7286 | 2.9057 | 5.0941 | 4.5620 | 7.6987 | 3.6468 |
| Std | 0.0108 | 0.0107 | 0.0086 | 0.0852 | 2.6354 | 2.8098 | 4.8009 | 6.4211 | 5.8353 | 2.9836 |

## B. Method

### B.1 Geo-Initializer

The multi-modality of the redundant torsions is shown in Figure. 7(c), with ‘N-C*α*-C-N’ and ‘C-N-C*α*-C’ demonstrating 3 to 4 peeks in the estimated distributions. In this mean, we select *K* = 4 in the von Mises mixtures. Besides, the encoders mapping the atom-level graph (𝒱_*at*_; ℰ_*at*_) is a three-layer GAT in implementation. Since the edge sets are composed of the bonds, the inter-chain messages cannot be passed. Therefore, the Geo-Initializer focuses more on the single-chain structure, leading the RMSDs to high values because of insufficient contextual conditions.

### B.2 Dynamic Weights

The weight of *Loss*_geo_ as *α*_3_ is changed gradually with exponential moving average updating (EMA). For the *α*_3_ in *t*-th iteration, it reads

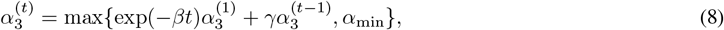

In which *α*_min_ is set as the minimum bond of the weight, *t* is the training iteration, *γ* and *β* are two decayed coefficients, which are hyper-parameters, set as 0.999 and 0.9999 in practice.

## C. Experiments

### C.1 Hyper-parameters

#### Heterogeneous residue-level encoder

is parameterized as 9 layers of heterogeneous GNNs. In each layer, the MLP is constructed by ‘`Linear + SiLu + Linear`’, with `Dropout` probability equaling 0.1 to avoid over-fitness. The `embedding dim` is set as 128.

#### Equivariant atom-level interaction layers

are composed of a 3-layers GAT. For each, the function *f* is parameterized by a three-layer MLP.

#### Training weights

*α*_1_, *α*_2_ are set as 1.0 and 0.8 respctively for co-design models. *α*_3_ is set as 0.4 in GeoAB-R and dynamic as Appendix. B.2 discussed. For ΔΔ*G* prediction, *α*_1_ is set as 1.0, and *α*_2_ and *α*_3_ are set as 0.4, because we hope the model can focus more on predicting accuracy and perceiving the structures gradually.

#### Training parameters

The learning rate *lr* is 5*e −* 4. In all training, the max training epoch is 20. `LambdaLR` schedule is used, with `lr lambda` is set as 0.95 *× lr*.

#### Epitope selection

In our main experiments on antibody CDR design, we select the 48 residues of the antigen closest to the antibody in terms of the C*α* distance as the epitope like (Jin et al., 2021).

### C.2 10-Cross Validation Results on SAbDab

Here we give the 10-CV evaluation results in SAbDab to show that GeoAB achieves the state-of-the-art performance in Table 8.

### C.3 Complete Analysis on Internal Geometries

We give the complete analysis of the generated internal geometries in Table. 9.

### C.4 ITA Results of GeoAB-R

Here we also conduct experiments on ITA with GeoAB-R, following the protocol of MEAN (Kong et al., 2023a). To ensure the expected generalizability, we select a total of 53 antibodies from SKEMPI V2.0 training set for affinity optimization and split SAbDab into training and validation sets in a ratio of 9:1 for pretraining the model. The results of ITA for antibody optimization are given in Table. 7, following the protocol of MEAN (Kong et al., 2023a).

**Table 7:** Results of ΔΔ*G* optimized by ITA with different methods.

| Region | $\Delta\Delta G$ - H3 |
| --- | --- |
| Random | 1.25 |
| RefineGNN | -3.41 |
| C-RefineGNN | -3.66 |
| Mean | -5.79 |
| C-DyMean | -6.56 |
| GeoAB-R | -7.28 |

**Table 8:** Results of 10-cross validation on different baselines. GeoAB-R and GeoAB-D reach overall the best performance.

| Methods | CDR-H1 |  | CDR-H2 |  | CDR-H3 |  |
| --- | --- | --- | --- | --- | --- | --- |
|  | AAR | RMSD | AAR | RMSD | AAR | RMSD |
| RefineGNN | 39.40 | 3.22 | 37.06 | 3.64 | 21.13 | 6.00 |
| C-RefineGNN | 33.19 | 3.25 | 33.53 | 3.69 | 18.88 | 6.22 |
| C-HERN | 47.36 | 1.45 | 41.45 | 1.20 | 28.47 | 2.64 |
| MEAN | 58.29 | 0.98 | 47.15 | 0.95 | 36.38 | 2.21 |
| C-DyMEAN | 61.13 | 0.89 | 53.14 | 0.87 | 37.09 | 2.09 |
| GeoAB-R | **64.87** | **0.82** | **56.09** | **0.74** | **37.58** | **1.94** |
| GeoAB-D | <u>64.03</u> | <u>0.84</u> | <u>54.98</u> | <u>0.75</u> | <u>37.09</u> | <u>1.97</u> |

**Table 9:**
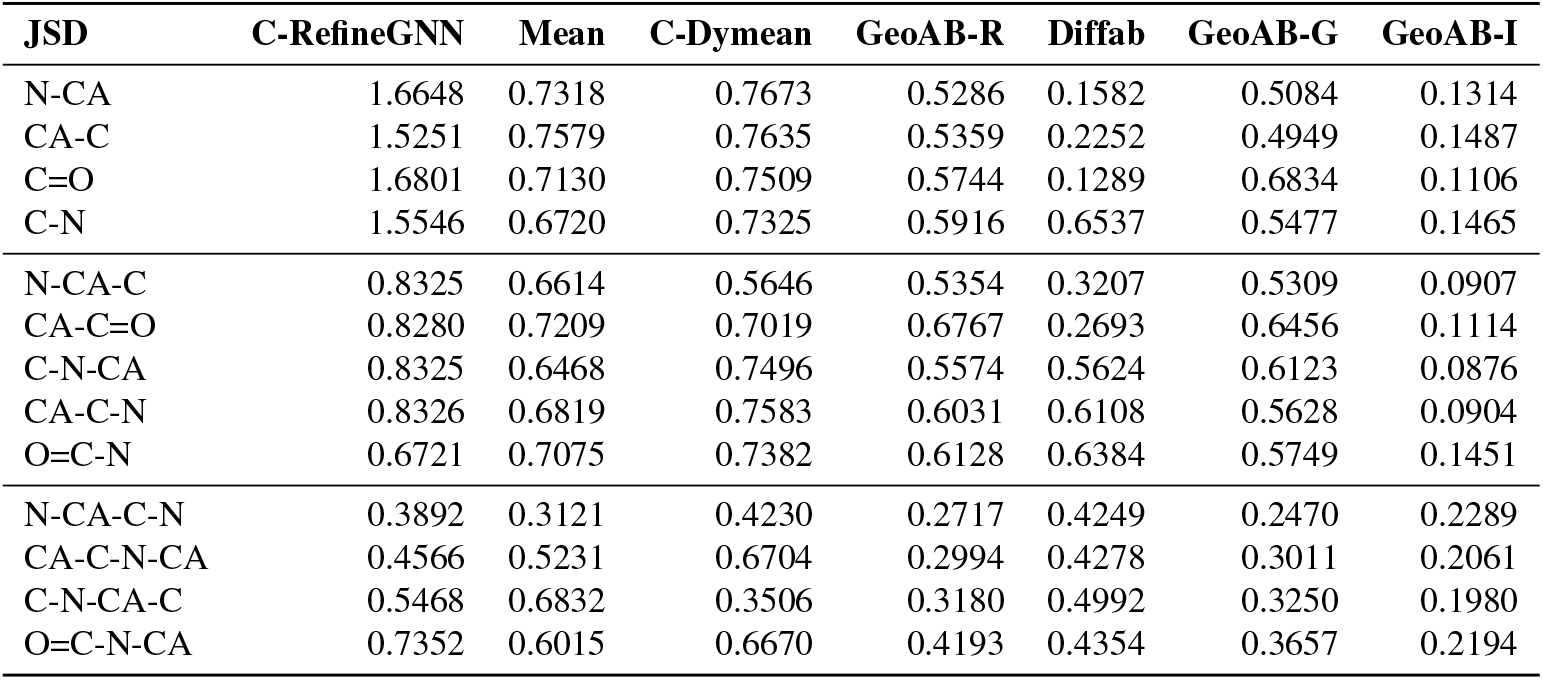
JSDs of the compared methods for different geometries *v*.*s*. the reference. GeoAB-I is the Geo-initializer.

| JSD | C-RefineGNN | Mean | C-Dymean | GeoAB-R | Diffab | GeoAB-G | GeoAB-I |
| --- | --- | --- | --- | --- | --- | --- | --- |
| N-CA | 1.6648 | 0.7318 | 0.7673 | 0.5286 | 0.1582 | 0.5084 | 0.1314 |
| CA-C | 1.5251 | 0.7579 | 0.7635 | 0.5359 | 0.2252 | 0.4949 | 0.1487 |
| C=O | 1.6801 | 0.7130 | 0.7509 | 0.5744 | 0.1289 | 0.6834 | 0.1106 |
| C-N | 1.5546 | 0.6720 | 0.7325 | 0.5916 | 0.6537 | 0.5477 | 0.1465 |
| N-CA-C | 0.8325 | 0.6614 | 0.5646 | 0.5354 | 0.3207 | 0.5309 | 0.0907 |
| CA-C=O | 0.8280 | 0.7209 | 0.7019 | 0.6767 | 0.2693 | 0.6456 | 0.1114 |
| C-N-CA | 0.8325 | 0.6468 | 0.7496 | 0.5574 | 0.5624 | 0.6123 | 0.0876 |
| CA-C-N | 0.8326 | 0.6819 | 0.7583 | 0.6031 | 0.6108 | 0.5628 | 0.0904 |
| O=C-N | 0.6721 | 0.7075 | 0.7382 | 0.6128 | 0.6384 | 0.5749 | 0.1451 |
| N-CA-C-N | 0.3892 | 0.3121 | 0.4230 | 0.2717 | 0.4249 | 0.2470 | 0.2289 |
| CA-C-N-CA | 0.4566 | 0.5231 | 0.6704 | 0.2994 | 0.4278 | 0.3011 | 0.2061 |
| C-N-CA-C | 0.5468 | 0.6832 | 0.3506 | 0.3180 | 0.4992 | 0.3250 | 0.1980 |
| O=C-N-CA | 0.7352 | 0.6015 | 0.6670 | 0.4193 | 0.4354 | 0.3657 | 0.2194 |

### C.5 Computational Comparison

We have conducted a comparison of our models with others on the CDR3 co-design task to address this concern. ‘Batch size’ is set to 8, layer number is 6 and the embed size is all set to 128 for a fair comparison. Table. 10. gives the complexity analysis. We can conclude that: In comparison to MEAN, which costs the least computational resources, our model has comparable GPU memory cost and training time, especially in GeoAB-R, because MEAN takes multi-head attention as its graph updating modules, while GeoAB-R uses a faster and simpler MLP-based heterogenous GNN. However, in the atom-updating layer, it is the key to increasing the training time because of a larger parameter number. For GeoAB-D, the initialization takes more time than Linear Initialization in GeoAB-R, which leads to more time in training and inference. However, since the initializer is pertained, so the trainable parameter number and GPU memory cost are of tiny significance compared with GeoAB-R. Note that DiffAB as a diffusion-based method, usually requires more training epochs and the iterative denoising process leads the inference time much longer than others.

**Table 10:**
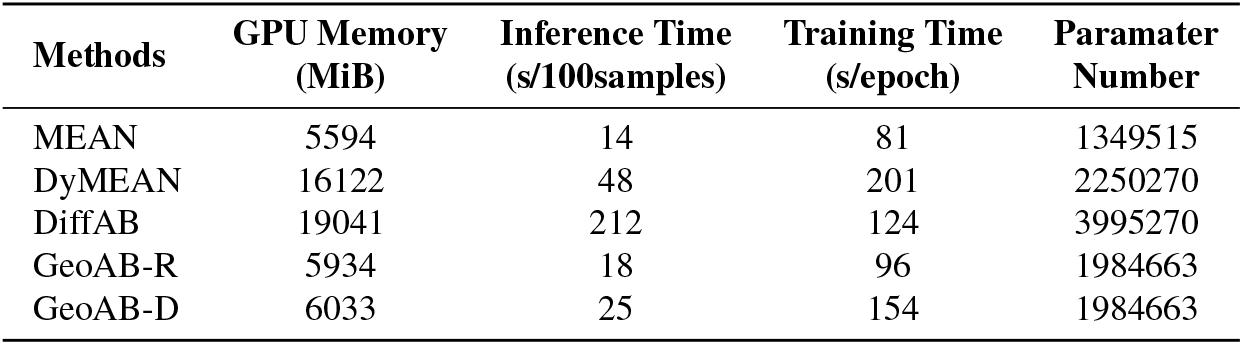
Complexity comparison of the included models.

| Methods | GPU Memory (MiB) | Inference Time (s/100samples) | Training Time (s/epoch) | Paramater Number |
| --- | --- | --- | --- | --- |
| MEAN | 5594 | 14 | 81 | 1349515 |
| DyMEAN | 16122 | 48 | 201 | 2250270 |
| DiffAB | 19041 | 212 | 124 | 3995270 |
| GeoAB-R | 5934 | 18 | 96 | 1984663 |
| GeoAB-D | 6033 | 25 | 154 | 1984663 |

## References

Adolf-Bryfogle, J., Kalyuzhniy, O., Kubitz, M., Weitzner, B. D., Hu, X., Adachi, Y., Schief, W. R., and Dunbrack Jr, R. L. Rosettaantibodydesign (rabd): A general framework for computational antibody design. PLoS computational biology, 14(4):e1006112, 2018.

Agnihotry, S., Pathak, R. K., Singh, D. B., Tiwari, A., and Hussain, I. Chapter 11 - protein structure prediction. In Singh, D. B. and Pathak, R. K. (eds.), Bioinformatics, pp. 177–188. Academic Press, 2022. ISBN 978-0-323-89775-4. doi: 10.1016/B978-0-323-89775-4.00023-7. URL https://www.sciencedirect.com/science/article/pii/B9780323897754000237.

Akbar, R., Robert, P. A., Weber, C. R., Widrich, M., Frank, R., Pavlović, M., Scheffer, L., Chernigovskaya, M., Snapkov, I., Slabodkin, A., et al. In silico proof of principle of machine learning-based antibody design at unconstrained scale. In Mabs, volume 14, pp. 2031482. Taylor & Francis, 2022.

Alford, R. F., Leaver-Fay, A., Jeliazkov, J. R., O’Meara, M. J., DiMaio, F. P., Park, H., Shapovalov, M. V., Renfrew, P. D., Mulligan, V. K., Kappel, K., et al. The rosetta all-atom energy function for macromolecular modeling and design. Journal of chemical theory and computation, 13 (6):3031–3048, 2017.

Alley, E. C., Khimulya, G., Biswas, S., AlQuraishi, M., and Church, G. M. Unified rational protein engineering with sequence-based deep representation learning. Nature methods, 16(12):1315–1322, 2019.

Basu, K., Green, E. M., Cheng, Y., and Craik, C. S. Why recombinant antibodies—benefits and applications. Current opinion in biotechnology, 60:153–158, 2019.

Cacabelos, R., Cacabelos, P., and Torrellas, C. Personalized medicine of alzheimer’s disease. Handbook of Pharmacogenomics and Stratified Medicine, pp. 563 – 615, 2014. URL https://api.semanticscholar.org/CorpusID:81213389.

Cai, H., Zhang, Z., Wang, M., Zhong, B., Wu, Y., Ying, T., and Tang, J. Pretrainable geometric graph neural network for antibody affinity maturation. bioRxiv, 2023. doi: 10.1101/2023.08.10.552845. URL https://www.biorxiv.org/content/early/2023/08/11/2023.08.10.552845.

Dauparas, J., Anishchenko, I. V., Bennett, N. R., Bai, H., Ragotte, R. J., Milles, L. F., Wicky, B. I. M., Courbet, A., de Haas, R. J., Bethel, N. P., Leung, P. J. Y., Huddy, T. F., Pellock, S. J., Tischer, D. K., Chan, F., Koepnick, B., Nguyen, H. A., Kang, A., Sankaran, B., Bera, A. K., King, N. P., and Baker, D. Robust deep learning based protein sequence design using proteinmpnn. Science (New York, N.Y.), 378:49 – 56, 2022. URL https://api.semanticscholar.org/CorpusID:249400681.

Delgado, J., Radusky, L. G., Cianferoni, D., and Serrano, L. Foldx 5.0: working with rna, small molecules and a new graphical interface. Bioinformatics, 35:4168 – 4169, 2019. URL https://api.semanticscholar.org/CorpusID:78094490.

Dunbar, J., Krawczyk, K., Leem, J., Baker, T., Fuchs, A., Georges, G., Shi, J., and Deane, C. M. Sabdab: the structural antibody database. Nucleic acids research, 42 (D1):D1140–D1146, 2014.

Gao, K., Wu, L., Zhu, J., Peng, T., Xia, Y., He, L., Xie, S., Qin, T., Liu, H., He, K., and Liu, T.-Y. Pretraining antibody language models for antigen-specific computational antibody design. In Proceedings of the 29th ACM SIGKDD Conference on Knowledge Discovery and Data Mining, KDD ‘23, pp. 506–517, New York, NY, USA, 2023. Association for Computing Machinery. ISBN 9798400701030. doi: 10.1145/3580305.3599468. URL 10.1145/3580305.3599468.

Geiger, M. and Smidt, T. e3nn: Euclidean neural networks, 2022.

Hsu, C., Verkuil, R., Liu, J., Lin, Z., Hie, B. L., Sercu, T., Lerer, A., and Rives, A. Learning inverse folding from millions of predicted structures. bioRxiv, 2022. URL https://api.semanticscholar.org/CorpusID:248151599.

Huang, W., Han, J., Rong, Y., Xu, T., Sun, F., and Huang, J. Equivariant graph mechanics networks with constraints. arXiv preprint arXiv:2203.06442, 2022.

Huang, Y., Li, S., Su, J., Wu, L., Zhang, O., Lin, H., Qi, J., Liu, Z., Gao, Z., Liu, Y., Zheng, J., and Li, S. Z. Protein 3d graph structure learning for robust structure-based protein property prediction. ArXiv, abs/2310.11466, 2023. URL https://api.semanticscholar.org/CorpusID:264288981.

Jankauskaitė, J., Jiménez-García, B., Dapkūnas, J., Fernández-Recio, J., and Moal, I. H. Skempi 2.0: an updated benchmark of changes in protein–protein binding energy, kinetics and thermodynamics upon mutation. Bioinformatics, 35(3):462–469, 2019.

Jin, W., Wohlwend, J., Barzilay, R., and Jaakkola, T. Iterative refinement graph neural network for antibody sequence-structure co-design. arXiv preprint arXiv:2110.04624, 2021.

Jin, W., Barzilay, R., and Jaakkola, T. Antibody-antigen docking and design via hierarchical structure refinement. In International Conference on Machine Learning, pp. 10217–10227. PMLR, 2022.

Jin, W., Sarkizova, S., Chen, X., Hacohen, N., and Uhler, C. Unsupervised protein-ligand binding energy prediction via neural euler’s rotation equation, 2023.

Jing, B., Corso, G., Chang, J., Barzilay, R., and Jaakkola, T. Torsional diffusion for molecular conformer generation, 2023.

Joosten, R. P., Long, F., Murshudov, G. N., and Perrakis, A. The pdb redo server for macromolecular structure model optimization. IUCrJ, 1:213 – 220, 2014. URL https://api.semanticscholar.org/CorpusID:10844981.

Jumper, J. M., Evans, R., Pritzel, A., Green, T., Figurnov, M., Ronneberger, O., Tunyasuvunakool, K., Bates, R., Zídek, A., Potapenko, A., Bridgland, A., Meyer, C., Kohl, S. A. A., Ballard, A., Cowie, A., Romera-Paredes, B., Nikolov, S., Jain, R., Adler, J., Back, T., Petersen, S., Reiman, D. A., Clancy, E., Zielinski, M., Steinegger, M., Pacholska, M., Berghammer, T., Bodenstein, S., Silver, D., Vinyals, O., Senior, A. W., Kavukcuoglu, K., Kohli, P., and Hassabis, D. Highly accurate protein structure prediction with alphafold. Nature, 596:583 – 589, 2021. URL https://api.semanticscholar.org/CorpusID:235959867.

Kong, X., Huang, W., and Liu, Y. Conditional antibody design as 3d equivariant graph translation. In The Eleventh International Conference on Learning Representations, 2023a. URL https://openreview.net/forum?id=LFHFQbjxIiP.

Kong, X., Huang, W., and Liu, Y. End-to-end full-atom antibody design, 2023b.

Lapidoth, G. D., Baran, D., Pszolla, G. M., Norn, C., Alon, A., Tyka, M. D., and Fleishman, S. J. Abdesign: A n algorithm for combinatorial backbone design guided by natural conformations and sequences. Proteins: Structure, Function, and Bioinformatics, 83(8):1385–1406, 2015.

Lefranc, M.-P., Pommié, C., Ruiz, M., Giudicelli, V., Foulquier, E., Truong, L., Thouvenin-Contet, V., and Lefranc, G. Imgt unique numbering for immunoglobulin and t cell receptor variable domains and ig superfamily v-like domains. Developmental & Comparative Immunology, 27(1):55–77, 2003.

Lei, R., Garcia, A. H., Tan, T. J. C., Teo, Q. W., Wang, Y., Zhang, X., Luo, S., Nair, S. K., Peng, J., and Wu, N. C. Mutational fitness landscape of human influenza h3n2 neuraminidase. Cell reports, 42:111951 – 111951, 2023. URL https://api.semanticscholar.org/CorpusID:255718376.

Leman, J. K., Weitzner, B. D., Lewis, S. M., Adolf-Bryfogle, J., Alam, N., Alford, R. F., Aprahamian, M. L., Baker, D., Barlow, K. A., Barth, P., Basanta, B., Bender, B. J., Blacklock, K. M., Bonet, J., Boyken, S. E., Bradley, P., Bystroff, C., Conway, P., Cooper, S., Correia, B. E., Coventry, B., Das, R., de Jong, R. M., DiMaio, F., Dsilva, L., Dunbrack, R. L., Ford, A., Frenz, B., Fu, D. Y., Geniesse, C., Goldschmidt, L., Gowthaman, R., Gray, J. J., Gront, D., Guffy, S. L., Horowitz, S., Huang, P.-S., Huber, T., Jacobs, T. M., Jeliazkov, J. R., Johnson, D. K., Kappel, K., Karanicolas, J., Khakzad, H., Khar, K. R., Khare, S. D., Khatib, F., Khramushin, A., King, I. C., Kleffner, R., Koepnick, B., Kortemme, T., Kuenze, G., Kuhlman, B., Kuroda, D., Labonte, J. W., Lai, J. K., Lapidoth, G., Leaver-Fay, A., Lindert, S., Linsky, T. W., London, N., Lubin, J. H., Lyskov, S., Maguire, J. B., Malmström, L., Marcos, E. S., Marcu, O., Marze, N. A., Meiler, J., Moretti, R., Mulligan, V. K., Nerli, S., Norn, C. H., ó ‘Conchuír, S., Ollikainen, N., Ovchinnikov, S., Pacella, M. S., Pan, X., Park, H., Pavlovicz, R. E., Pethe, M. A., Pierce, B. G., Pilla, K. B., Raveh, B., Renfrew, P. D., Burman, S. S. R., Rubenstein, A. B., Sauer, M., Scheck, A., Schief, W. R., Schueler-Furman, O., Sedan, Y., Sevy, A. M., Sgourakis, N. G., Shi, L., Siegel, J. B., Silva, D., Smith, S. T., Song, Y., Stein, A., Szegedy, M. A., Teets, F. D., Thyme, S. B., Wang, R. Y.-R., Watkins, A. M., Zimmerman, L., and Bonneau, R. Macromolecular modeling and design in rosetta: recent methods and frameworks. Nature Methods, 17:665 – 680, 2020. URL https://api.semanticscholar.org/CorpusID:219170571.

Li, T., Pantazes, R. J., and Maranas, C. D. Optmaven–a new framework for the de novo design of antibody variable region models targeting specific antigen epitopes. PloS one, 9(8):e105954, 2014.

Lieberman, M. A. Comprar marks’ essentials of medical biochemistry, 2a ed. a clinical approach — michael lieberman — 9781451190069 — lippincott williams & wilkins. 2014. URL https://api.semanticscholar.org/CorpusID:164543399.

Lin, H., Huang, Y., Liu, M., Li, X., Ji, S., and Li, S. Z. Diffbp: Generative diffusion of 3d molecules for target protein binding, 2022.

Lin, H., Huang, Y., Zhang, O., Wu, L., Li, S., Chen, Z., and Li, S. Z. Functional-group-based diffusion for pocket-specific molecule generation and elaboration, 2023.

Liu, G., Zeng, H., Mueller, J., Carter, B., Wang, Z., Schilz, J., Horny, G., Birnbaum, M. E., Ewert, S., and Gifford, D. K. Antibody complementarity determining region design using high-capacity machine learning. Bioinformatics, 36(7):2126–2133, 2020.

Liu, M., Luo, Y., Wang, L., Xie, Y., Yuan, H., Gui, S., Xu, Z., Yu, H., Zhang, J., Liu, Y., Yan, K., Oztekin, B., Liu, H., Zhang, X., Fu, C., and Ji, S. Dig: A turnkey library for diving into graph deep learning research. ArXiv, abs/2103.12608, 2021. URL https://api.semanticscholar.org/CorpusID:232320529.

Luo, S., Su, Y., Peng, X., Wang, S., Peng, J., and Ma, J. Antigen-specific antibody design and optimization with diffusion-based generative models for protein structures. bioRxiv, 2022. URL https://api.semanticscholar.org/CorpusID:250534060.

Luo, S., Su, Y., Wu, Z., Su, C., Peng, J., and Ma, J. Rotamer density estimator is an unsupervised learner of the effect of mutations on protein-protein interaction. In The Eleventh International Conference on Learning Representations, 2023. URL https://openreview.net/forum?id=_X9Yl1K2mD.

MacArthur, M. W. and Thornton, J. M. Deviations from planarity of the peptide bond in peptides and proteins. Journal of molecular biology, 264 5:1180– 95, 1996. URL https://api.semanticscholar.org/CorpusID:2831927.

Maier, J. A., Martinez, C., Kasavajhala, K., Wickstrom, L., Hauser, K. E., and Simmerling, C. ff14sb: Improving the accuracy of protein side chain and backbone parameters from ff99sb. Journal of Chemical Theory and Computation, 11(8):3696–3713, 2015. doi: 10.1021/acs.jctc.5b00255. URL 10.1021/acs.jctc.5b00255. PMID: 26574453.

Meier, J., Rao, R., Verkuil, R., Liu, J., Sercu, T., and Rives, A. Language models enable zero-shot prediction of the effects of mutations on protein function. bioRxiv, 2021. doi: 10.1101/2021.07.09.450648. URL https://www.biorxiv.org/content/early/2021/07/10/2021.07.09.450648.

Min-yi, S. and Andrej, S. Statistical potential for assessment and prediction of protein structures. Protein Sci., 15(11): 2507–24, 2006.

Park, H., Bradley, P., Greisen, P. J., Liu, Y., Mulligan, V. K., Kim, D. E., Baker, D., and DiMaio, F. Simultaneous optimization of biomolecular energy functions on features from small molecules and macromolecules. Journal of Chemical Theory and Computation, 12(12):6201–6212, 2016. doi: 10.1021/acs.jctc.6b00819. URL 10.1021/acs.jctc.6b00819. xPMID: 27766851.

Parsons, J., Holmes, J., Rojas, J., Tsai, J., and Strauss, C. Practical conversion from torsion space to cartesian space for in silico protein synthesis. Journal of computational chemistry, 26:1063–8, 07 2005. doi: 10.1002/jcc.20237.

Robinson, S. W., Afzal, A. M., and Leader, D. P. Chapter 13 - bioinformatics: Concepts, methods, and data. In Padmanabhan, S. (ed.), Handbook of Pharmacogenomics and Stratified Medicine, pp. 259–287. Academic Press, San Diego, 2014. ISBN 978-0-12-386882-4. doi: 10.1016/B978-0-12-386882-4.00013-X. URL https://www.sciencedirect.com/science/article/pii/B978012386882400013X.

Saka, K., Kakuzaki, T., Metsugi, S., Kashiwagi, D., Yoshida, K., Wada, M., Tsunoda, H., and Teramoto, R. Antibody design using lstm based deep generative model from phage display library for affinity maturation. Scientific reports, 11(1):1–13, 2021.

Satorras, V. G., Hoogeboom, E., and Welling, M. E (n) equivariant graph neural networks. In International Conference on Machine Learning, pp. 9323–9332. PMLR, 2021.

Shan, S., Luo, S., Yang, Z., Hong, J., Su, Y., Ding, F., Fu, L., Li, C., Chen, P., Ma, J., et al. Deep learning guided optimization of human antibody against sars-cov-2 variants with broad neutralization. Proceedings of the National Academy of Sciences, 119(11):e2122954119, 2022.

Steinegger, M. and Söding, J. Mmseqs2 enables sensitive protein sequence searching for the analysis of massive data sets. Nature biotechnology, 35(11):1026–1028, 2017.

Sulea, T., Hussack, G., Ryan, S., Tanha, J., and Purisima, E. O. Application of assisted design of antibody and protein therapeutics (adapt) improves efficacy of a clostridium difficile toxin a single-domain antibody. Scientific Report, 8, 2018.

Swanson, K., Williams, J., and Jonas, E. Von mises mixture distributions for molecular conformation generation, 2023.

Tan, C., Zhang, Y., Gao, Z., Hu, B., Li, S., Liu, Z., and Li, S. Z. Hierarchical data-efficient representation learning for tertiary structure-based rna design. In The Twelfth International Conference on Learning Representations, 2023.

Tan, C., Gao, Z., Wu, L., Xia, J., Zheng, J., Yang, X., Liu, Y., Hu, B., and Li, S. Z. Cross-gate mlp with protein complex invariant embedding is a one-shot antibody designer. In Proceedings of the AAAI Conference on Artificial Intelligence, volume 38, pp. 15222–15230, 2024.

Veličković, P., Cucurull, G., Casanova, A., Romero, A., Liò, P., and Bengio, Y. Graph attention networks, 2018.

Villar, S., Hogg, D. W., Storey-Fisher, K., Yao, W., and Blum-Smith, B. Scalars are universal: Equivariant machine learning, structured like classical physics, 2023.

Wu, F. and Li, S. Z. A hierarchical training paradigm for antibody structure-sequence co-design, 2023.

Wu, K. E., Yang, K. K., van den Berg, R., Zou, J., Lu, A. X., and Amini, A. P. Protein structure generation via folding diffusion. ArXiv, abs/2209.15611, 2022. URL https://api.semanticscholar.org/CorpusID:252668551.

Wu, L., Lin, H., Gao, Z., Tan, C., and Stan. Z. Li. Self-supervised learning on graphs: Contrastive, generative, or predictive. IEEE Transactions on Knowledge and Data Engineering, 35:4216–4235, 2021. URL https://api.semanticscholar.org/CorpusID:238215156.

Wu, L., Lin, H., Huang, Y., and Li, S. Z. Quantifying the knowledge in gnns for reliable distillation into mlps. In International Conference on Machine Learning, 2023. URL https://api.semanticscholar.org/CorpusID:259129782.

Wu, L., Tian, Y., Huang, Y., Li, S., Lin, H., Chawla, N., and Li, S. Z. Mape-ppi: Towards effective and efficient protein-protein interaction prediction via microenvironment-aware protein embedding. ArXiv, abs/2402.14391, 2024. URL https://api.semanticscholar.org/CorpusID:267782631.

Yang, K., Jin, W., Swanson, K., Barzilay, R., and Jaakkola, T. Improving molecular design by stochastic iterative target augmentation. In International Conference on Machine Learning, pp. 10716–10726. PMLR, 2020.

Yang, K. K., Zanichelli, N., and Yeh, H. Masked inverse folding with sequence transfer for protein representation learning. bioRxiv, 2023. URL https://api.semanticscholar.org/CorpusID:249241961.

